# DeepCor: Denoising fMRI Data with Contrastive Autoencoders

**DOI:** 10.1101/2023.10.31.565011

**Authors:** Yu Zhu, Aidas Aglinskas, Stefano Anzellotti

**Affiliations:** Department of Psychology and Neuroscience Boston College, Boston, MA 02467

## Abstract

Functional magnetic resonance imaging (fMRI) is widely used in neuroscience research. FMRI data is noisy; improving denoising methods could lead to novel discoveries. Here, we introduce and evaluate a denoising method (DeepCor) which utilizes deep generative models to disentangle and remove noise. DeepCor outperforms CompCor (a state-of-the art denoising approach) on a variety of simulated datasets. In addition, DeepCor enhances differences in connectivity between brain networks in real datasets.

---

FMRI is widely used in cognitive and clinical research [1, 2]. Improving fMRI analysis can support progress in multiple fields. FMRI measurements are affected by a variety of noise sources [3]. Noise can conceal phenomena of interest. Thus, advances in denoising could reveal phenomena that are currently hidden. Removing noise effectively is particularly important when data is scarce; for example, when the goal is to characterize brain activity in individual participants [4].Therefore, progress in denoising is important for the understanding of individual variation and for personalized psychiatry [5, 6].

A popular method – CompCor [7] – extracts responses from regions-of-no-interest (RONI, such as cerebrospinal fluid), and regresses them out from responses in regions-of interest (ROI, such as gray matter). An advantage of this strategy is that measurements in the RONI can capture how changes at the level of the noise sources (e.g. head movements) nonlinearly impact measurements at the voxel level (see [8]). A limitation is that this strategy assumes measurements to result from a linear summation of signal and noise. Instead, signal and noise could interact nonlinearly.

To overcome this limitation, the field is shifting towards denoising approaches based on deep learning ([9, 10, 11, 12]). While initial results are promising, some of these approaches are only applicable to task-based experiments [9], and others require large numbers of participants [11]. In this study, we introduce a novel approach – DeepCor – that can be applied to task-based experiments as well as resting-state data, and can be trained within individual participants. DeepCor takes advantage of probabilistic generative models that have been effective in other recent applications [13, 14].

DeepCor uses voxels from an ROI, where measurements contain both neural signal and noise, and from an RONI, where measurements contain only noise (Figure 1a-b). A contrastive variational autoencoder (CVAE) [13] is trained to disentangle features unique to timecourses extracted from the ROI from features that are also present in the RONI. (Figure 1c). Finally, denoised timecourses are generated by setting the features that are also present in the RONI to zero, and using the CVAE’s decoder to produce the denoised data (Figure 1d).

**Fig. 1.**
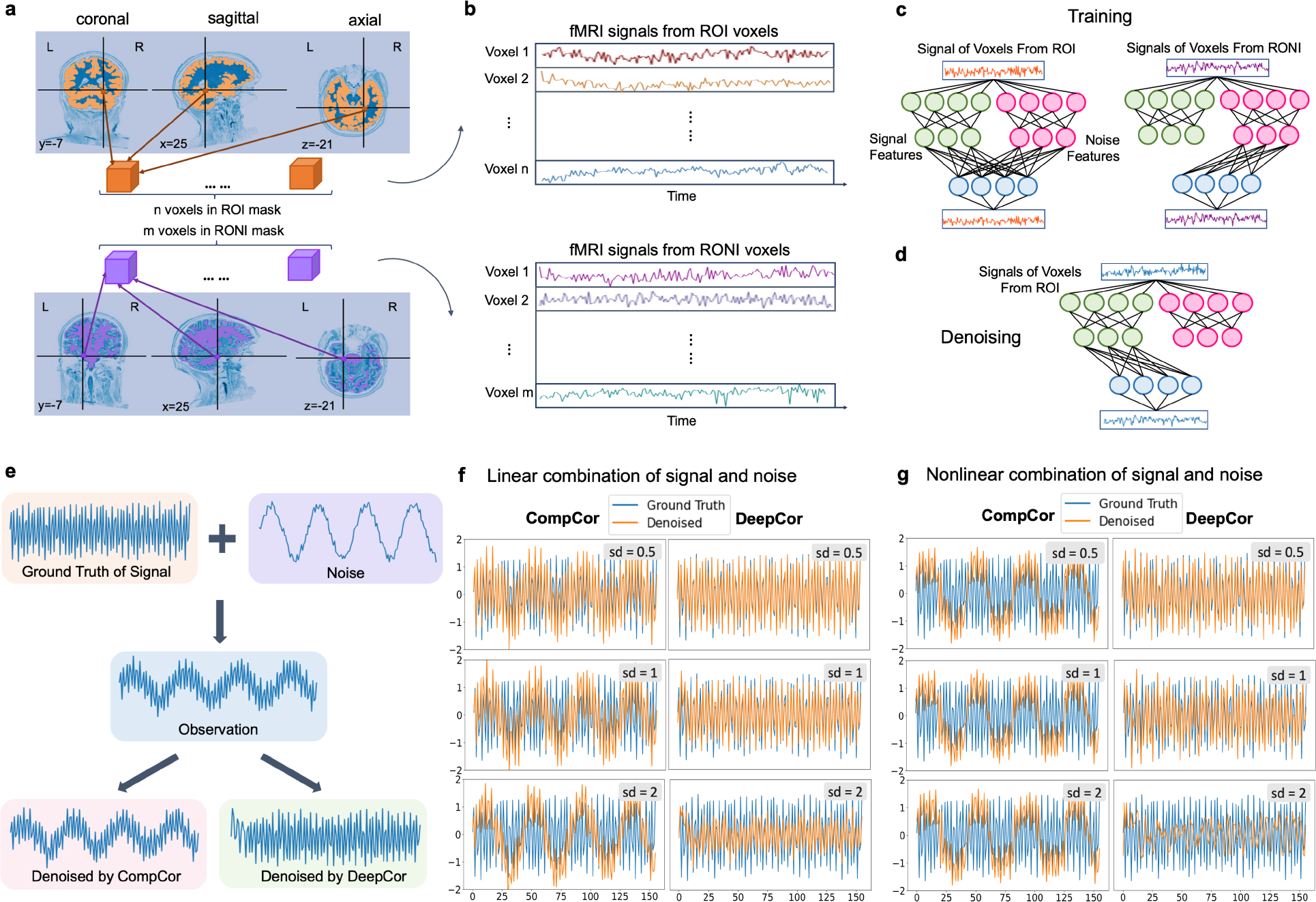
a) Identifying voxels from ROI and RONI masks. b) Extracting fMRI timecourses. c) Training the CVAE model. d) Denoising. e) A example of simuated data. f-g) Examples of denoised data compared to the ground truth. Different rows correspond to different amounts of noise.

In fMRI experiments the ground truth of the neural signal is not known, therefore we started by testing DeepCor using simulated data. In a first set of simulations, we created the signal and noise with trigonometric functions (Figure 1e). In order to evaluate DeepCor’s ability to denoise data under different conditions, we tested two ways of combining the ground truth with the noise: with a linear function of the ground truth and the noise (their sum), and with a nonlinear function (the ground truth plus the cubic root of the noise). We then denoised the dataset using both CompCor [7] and DeepCor.

Data denoised with DeepCor matched the ground truth more closely (as measured with R-squared), achieving a 17% improvement over aCompCor (*w*(9999) = 50004996.0, *p <* 0.001) when the ground truth and noise were combined linearly, and a 56% improvement (*w*(9999) = 49963547.0, *p <* 0.001) when the ground truth and noise were combined nonlinearly (see Figure 1f-g). We then tested the impact of the amount of noise on the results (Table 4). As expected, increasing the amount of noise reduced the correlations between the denoised data and the ground truth overall, but DeepCor consistently achieved higher R-squared values than CompCor.

In order to evaluate DeepCor using more realistic simulated data, we used BrainIAK [15, 16, 17] (see https://brainiak.org/examples/fmrisim_multivariate_example.html, Figure 2a). We generated noise utilizing parameters derived from empirical fMRI data [18] (Figure 2b). The ground truth of the signal was simulated as responses to two distinct event types (Figure 2c), associated with different beta-values in different ROI voxels. Simulated ROI data were created by summing the noise and the ground truth of the signal. The dataset was denoised with DeepCor and CompCor. DeepCor yielded a R-squared value improvement of 402% compared to CompCor (*w*(15478) = 119807460.0, *p <* 0.001, Figure 2d-e). These results demonstrate that even for more realistic datasets, DeepCor outperforms CompCor, yielding denoised data that are more similar to the ground truth.

**Fig. 2.**
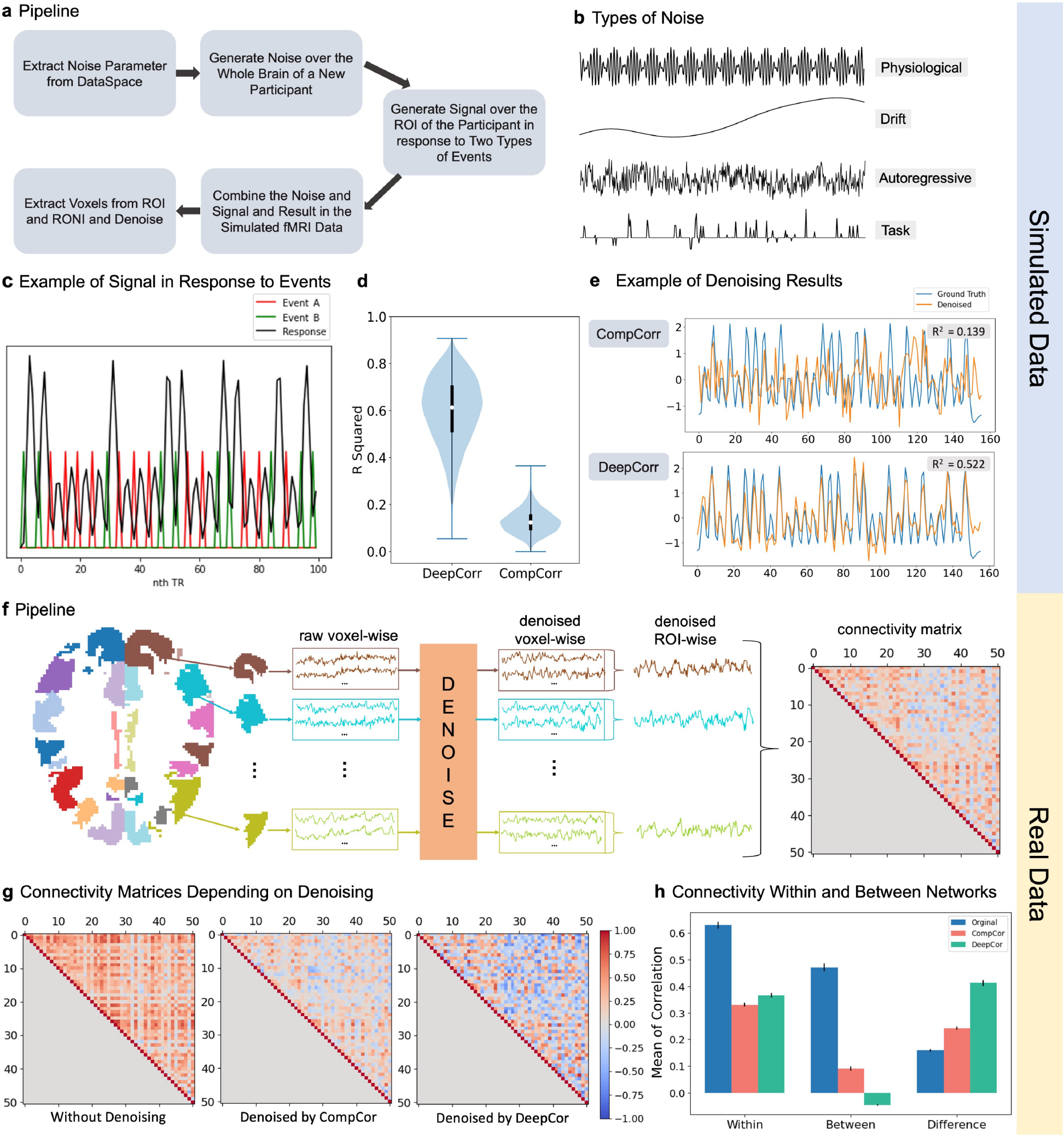
a) BrainIAK simulation pipeline. b) Noise types. c) Signal simulation. d) Correspondence between denoised timecourses and the ground truth. e) Examples of denoised timeseries compared to the ground truth. f) Denoising of real data in connectivity-defined regions of interest. g) Example connectivity matrices before and after denoising. h) Impact of denoising on functional connectivity within and between networks.

Having established that DeepCor outperforms CompCor across a variety of simulations, we set out to study its impact on real fMRI data. One application of fMRI involves the analysis of the interactions between brain regions: functional connectivity. Based on patterns of connectivity, cortex can be subdivided into 51 regions (Figure 2f) and 7 networks [19], such that regions within the same network have high functional connectivity with each other. This observation offers an opportunity to evaluate the impact of DeepCor on real fMRI data. In fact, noise in fMRI measurements can affect the whole brain, and can lead to spurious functional connectivity even between brain regions that do not have any functional coupling. Considering this, we predicted that 1) CompCor and DeepCor would reduce the functional connectivity between brain regions in different functional networks compared to the raw data (this serves as a “sanity check”), and 2) DeepCor would lead to a grater difference in connectivity between pairs of regions within the same network and pairs of regions in different networks compared to CompCor, by enhancing connectivity between regions within the same network, and decreasing connectivity between regions in different networks. In order to test this, we analyzed resting state fMRI data from 200 participants in the ABIDE dataset [20]. We computed connectivity matrices between the 51 regions for the raw data, the data denoised with CompCor, and the data denoised with DeepCor (Figure 2g). Results were in line with the expectations: the difference in connectivity for regions within the same network vs regions in different networks was significantly larger in data denoised with DeepCor (*w*(199) = 19977.0, *p <* 0.001) (Figure 2h).

In conclusion, DeepCor outperformed CompCor in all tests. The results demonstrate the effectiveness of DeepCor for noise reduction, across simulated and real data. By treating individual voxels as data points, DeepCor obtains sufficient data to be trained independently within each individual participant even in relatively short fMRI scans, therefore it can be flexibly applied to studies of individual differences. In addition, it does not require information about the experimental task, so it can be applied to both task-based experiments and resting state experiments.

## Method

### Overview of DeepCor

DeepCor is a method for denoising fMRI data based on probabilistic deep learning. DeepCor uses the responses in regions-of-no-interest (RONI) that contain only noise (e.g. cerebrospinal fluid and white matter) to identify and remove noise from regions-of-interest (ROI, e.g. gray matter). Using contrastive variational autoencoders (CVAEs, [13]), DeepCor identifies features that are in common to measurements in the ROI and the RONI and separates them from features that are unique to measurements in the ROI. Taking advantage of the fact that CVAEs are generative models, DeepCor performs denoising by setting to zero the features in common to both the ROI and the RONI, and reconstructing the data. Much of the noise, which is in common to the ROI and the RONI, is therefore removed. A similar logic has been adopted in previous work to disentangle disorder-specific variation from variation shared with the general population [14].

### DeepCor: Technical Details

#### Inputs

Consider a functional MRI dataset of dimensions *L*× *W*× *H*× *T*, where *L*× *W*× *H* is the number of voxels and *T* is the number of timepoints. We use a mask of the RONI to extract the timecourses of the *m* RONI voxels, and a mask of the ROI to extract the timecourses of the *n* ROI voxels. In this study, the ROI mask was generated by thresholding the probability map for gray matter obtained during segmentation with a *p >* 0.50 threshold, and the RONI mask was generated by thresholding the sum of the probability maps for white matter and cerebrospinal fluid with a *p >* 0.50 threshold. Each timecourse was normalized by subtracting its mean and dividing by its standard deviation.

#### Model structure

The model is an autoencoder consisting of two probabilistic encoders: 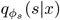, which will yield as output features of the signal, and 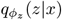, which will yield as output features of the noise. The measurement is then reconstructed with a decoder *f*_*θ*_(*s, z*) that takes as input the concatenated outputs of the two encoders. Each encoder consists of four 1D convolutional layers and two parallel fully-connected layers representing parameters for mean and standard deviation of the distribution, and the decoder is composed by one fully-connected layer connecting to the latent space and four 1D transposed convolutional layer followed by one final convolutional layer. Detailed hyperparameters are indicated in Tables 1-3.

**Table 1.** CVAE architecture for simulated dataset with trigonometric functions

| Block | Layer | Kernel | Stride | Padding | Out-Padding | Activation Function | Input Size | Output Size | Input Source |
| --- | --- | --- | --- | --- | --- | --- | --- | --- | --- |
| Encoder $q_{\phi_s}(s\vert x)$ | Conv1d(1) | 3 | 2 | 1 | - | BatchNorm, LeakyReLU | $1 \times 1 \times 156$ | $32 \times 1 \times 78$ | Input |
| | Conv1d(2) | 3 | 2 | 1 | - | BatchNorm, LeakyReLU | $32 \times 1 \times 78$ | $64 \times 1 \times 39$ | Conv1d(1) |
| | Conv1d(3) | 3 | 2 | 1 | - | BatchNorm, LeakyReLU | $64 \times 1 \times 39$ | $128 \times 1 \times 20$ | Conv1d(2) |
| | Conv1d(4) | 3 | 2 | 1 | - | BatchNorm, LeakyReLU | $128 \times 1 \times 20$ | $128 \times 1 \times 10$ | Conv1d(3) |
| | Flatten | - | - | - | - | - | $128 \times 1 \times 10$ | 1280 | Conv1d(4) |
| | FC $\mu_s$ | - | - | - | - | - | 1280 | 16 | Flatten |
| | FC $\sigma_s^2$ | - | - | - | - | - | 1280 | 16 | Flatten |
| | Reparametrize | - | - | - | - | - | 16 | 16 | FC $\mu_s$, FC $\sigma_s^2$ |
| Encoder $q_{\phi_n}(n\vert x)$ | Conv1d(1) | 3 | 2 | 1 | - | BatchNorm, LeakyReLU | $1 \times 1 \times 156$ | $32 \times 1 \times 78$ | Inputs |
| | Conv1d(2) | 3 | 2 | 1 | - | BatchNorm, LeakyReLU | $32 \times 1 \times 78$ | $64 \times 1 \times 39$ | Conv1d(1) |
| | Conv1d(3) | 3 | 2 | 1 | - | BatchNorm, LeakyReLU | $64 \times 1 \times 39$ | $128 \times 1 \times 20$ | Conv1d(2) |
| | Conv1d(4) | 3 | 2 | 1 | - | BatchNorm, LeakyReLU | $128 \times 1 \times 20$ | $128 \times 1 \times 10$ | Conv1d(3) |
| | Flatten | - | - | - | - | - | $128 \times 1 \times 10$ | 1280 | Conv1d(4) |
| | FC $\mu_n$ | - | - | - | - | - | 1280 | 16 | Flatten |
| | FC $\sigma_n^2$ | - | - | - | - | - | 1280 | 16 | Flatten |
| | Reparametrize | - | - | - | - | - | 16 | 16 | FC $\mu_n$, FC $\sigma_n^2$ |
| Latent Space | Concatenate | - | - | - | - | - | 16,16 | 32 | $q_{\phi_s}, q_{\phi_n}$ |
| Decoder $f_{\theta}(s, n)$ | FC | - | - | - | - | - | 32 | 1280 | Latent Space |
| | Reshape | - | - | - | - | - | 1280 | $128 \times 1 \times 10$ | FC |
| | ConvTrans1d(1) | 3 | 2 | 1 | 1 | BatchNorm, LeakyReLU | $128 \times 1 \times 10$ | $128 \times 1 \times 20$ | Reshape |
| | ConvTrans1d(2) | 3 | 2 | 1 | 1 | BatchNorm, LeakyReLU | $128 \times 1 \times 20$ | $64 \times 1 \times 40$ | ConvTrans1d(1) |
| | ConvTrans1d(3) | 3 | 2 | 1 | - | BatchNorm, LeakyReLU | $64 \times 1 \times 40$ | $32 \times 1 \times 79$ | ConvTrans1d(2) |
| | ConvTrans1d(4) | 3 | 2 | 1 | 1 | BatchNorm, LeakyReLU | $32 \times 1 \times 79$ | $32 \times 1 \times 158$ | ConvTrans1d(3) |
| | Conv1d | 3 | 1 | 1 | - | BatchNorm, LeakyReLU | $32 \times 1 \times 158$ | $1 \times 1 \times 156$ | ConvTrans1d(4) |

**Table 2.** CVAE architecture for realistic data simulation

| Block | Layer | Kernel | Stride | Padding | Out-Padding | Activation Function | Input Size | Output Size | Input Source |
| --- | --- | --- | --- | --- | --- | --- | --- | --- | --- |
| Encoder<br>$q_{\phi_s}(s\vert x)$ | Conv1d(1) | 3 | 2 | 1 | - | BatchNorm,<br>LeakyReLU | $1 \times 1 \times 156$ | $64 \times 1 \times 78$ | Input |
| | Conv1d(2) | 3 | 2 | 1 | - | BatchNorm,<br>LeakyReLU | $64 \times 1 \times 78$ | $128 \times 1 \times 39$ | Conv1d(1) |
| | Conv1d(3) | 3 | 2 | 1 | - | BatchNorm,<br>LeakyReLU | $128 \times 1 \times 39$ | $256 \times 1 \times 20$ | Conv1d(2) |
| | Conv1d(4) | 3 | 2 | 1 | - | BatchNorm,<br>LeakyReLU | $256 \times 1 \times 20$ | $256 \times 1 \times 10$ | Conv1d(3) |
| | Flatten | - | - | - | - | - | $256 \times 1 \times 10$ | 2560 | Conv1d(4) |
| | FC $\mu_s$ | - | - | - | - | - | 2560 | 8 | Flatten |
| | FC $\sigma_s^2$ | - | - | - | - | - | 2560 | 8 | Flatten |
| | Reparameterize | - | - | - | - | - | 8 | 8 | FC $\mu_s, \text{FC } \sigma_s^2$ |
| | Conv1d(2) | 3 | 2 | 1 | - | BatchNorm,<br>LeakyReLU | $64 \times 1 \times 78$ | $128 \times 1 \times 39$ | Conv1d(1) |
| | Conv1d(3) | 3 | 2 | 1 | - | BatchNorm,<br>LeakyReLU | $128 \times 1 \times 39$ | $256 \times 1 \times 20$ | Conv1d(2) |
| | Conv1d(4) | 3 | 2 | 1 | - | BatchNorm,<br>LeakyReLU | $256 \times 1 \times 20$ | $256 \times 1 \times 10$ | Conv1d(3) |
| | Flatten | - | - | - | - | - | $256 \times 1 \times 10$ | 2560 | Conv1d(4) |
| | FC $\mu_n$ | - | - | - | - | - | 2560 | 8 | Flatten |
| | FC $\sigma_n^2$ | - | - | - | - | - | 2560 | 8 | Flatten |
| | Reparameterize | - | - | - | - | - | 8 | 8 | FC $\mu_n, \text{FC } \sigma_n^2$ |
| Latent Space | Concatenate | - | - | - | - | - | 8,8 | 16 | $q_{\phi_s}, q_{\phi_n}$ |
| Decoder<br>$f_{\theta}(s, n)$ | FC | - | - | - | - | - | 16 | 2560 | Latent Space |
| | Reshape | - | - | - | - | - | 2560 | $256 \times 1 \times 10$ | FC |
| | ConvTrans1d(1) | 3 | 2 | 1 | 1 | BatchNorm,<br>LeakyReLU | $256 \times 1 \times 10$ | $256 \times 1 \times 20$ | Reshape |
| | ConvTrans1d(2) | 3 | 2 | 1 | 1 | BatchNorm,<br>LeakyReLU | $256 \times 1 \times 20$ | $128 \times 1 \times 40$ | ConvTrans1d(1) |
| | ConvTrans1d(3) | 3 | 2 | 1 | - | BatchNorm,<br>LeakyReLU | $128 \times 1 \times 40$ | $64 \times 1 \times 79$ | ConvTrans1d(2) |
| | ConvTrans1d(4) | 3 | 2 | 1 | 1 | BatchNorm,<br>LeakyReLU | $64 \times 1 \times 79$ | $64 \times 1 \times 158$ | ConvTrans1d(3) |
| | Conv1d | 3 | 1 | 1 | - | BatchNorm,<br>LeakyReLU | $64 \times 1 \times 158$ | $1 \times 1 \times 156$ | ConvTrans1d(4) |

**Table 3.**
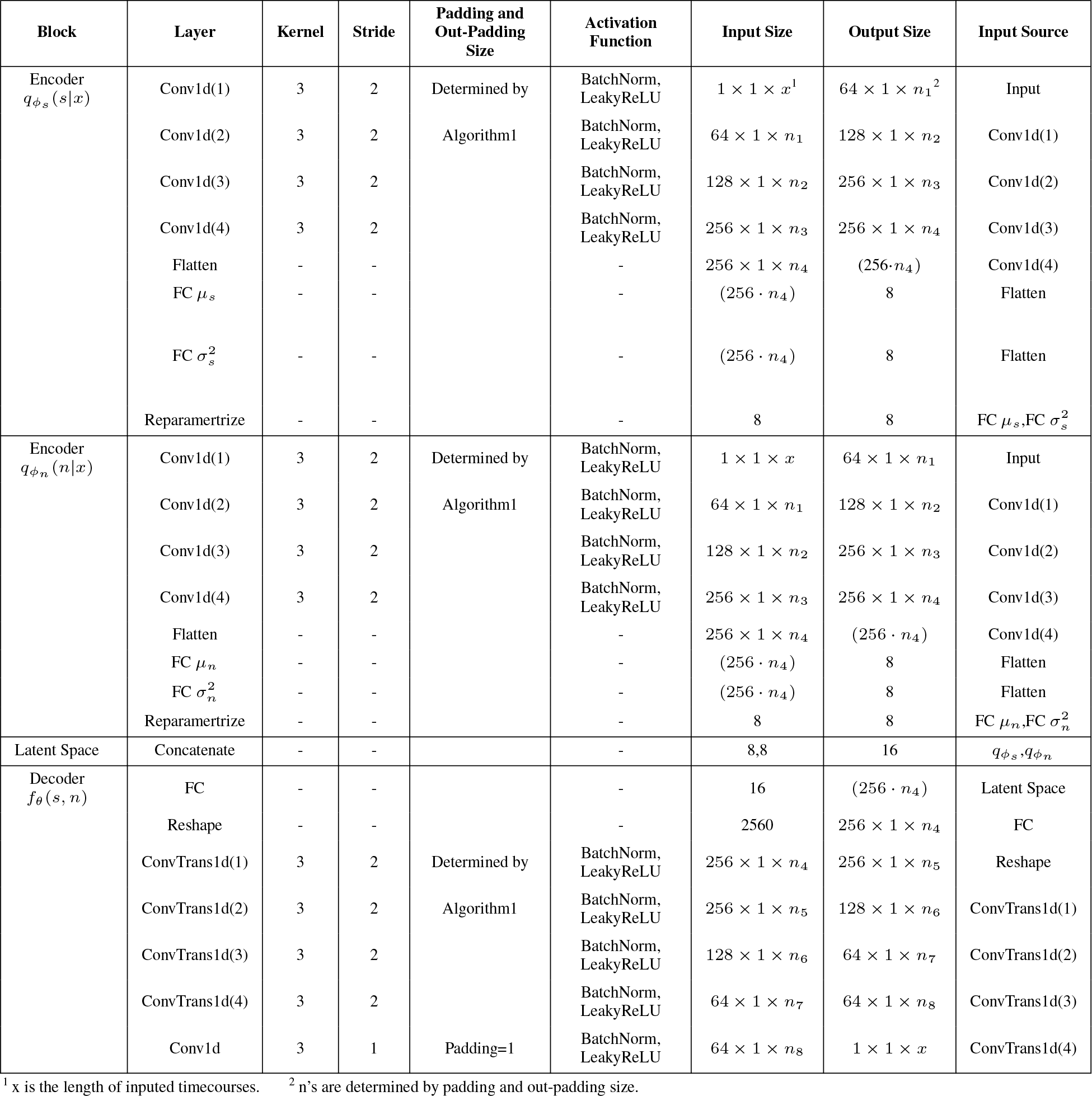
CVAE architecture for functional connectivity analysis

**Table 4.** Training Effiency of DeepCor and CompCor in Comparison

|  | Trigonometric Linear |  |  | Trigonometric Non-linear |  |  | BrainIAK |
| --- | --- | --- | --- | --- | --- | --- | --- |
| Standard deviation | 0.5 | 1.0 | 2.0 | 0.5 | 1.0 | 2.0 |  |
| DeepCor $R^2$ | 0.9631 | 0.9303 | 0.7891 | 0.9520 | 0.8378 | 0.4163 | 0.6090 |
| DeepCor $R^2$<br>5%-95% Percentile | (0.9456,<br>0.9763) | (0.9042,<br>0.9545) | (0.7110,<br>0.8577) | (0.9364,<br>0.9645) | (0.7985,<br>0.8793) | (0.3238,<br>0.5300) | (0.3835,<br>0.8172) |
| CompCor $R^2$ | 0.8232 | 0.5460 | 0.2434 | 0.8197 | 0.5382 | 0.2290 | 0.1213 |
| CompCor $R^2$<br>5%-95% Percentile | (0.7201,<br>0.9283) | (0.3919,<br>0.7489) | (0.1379,<br>0.4364) | (0.7758,<br>0.8955) | (0.4614,<br>0.7271) | (0.1729,<br>0.3771) | (0.0520,<br>0.2134) |
| Improvement <sup>1</sup> | 17% | 70% | 224% | 17% | 56% | 82% | 402% |

<sup>1</sup> DeepCor’s improvement over CompCor, which equals $(\frac{DeepCor R^2}{CompCor R^2} - 1) \times 100\%$ .

**Table 5.**
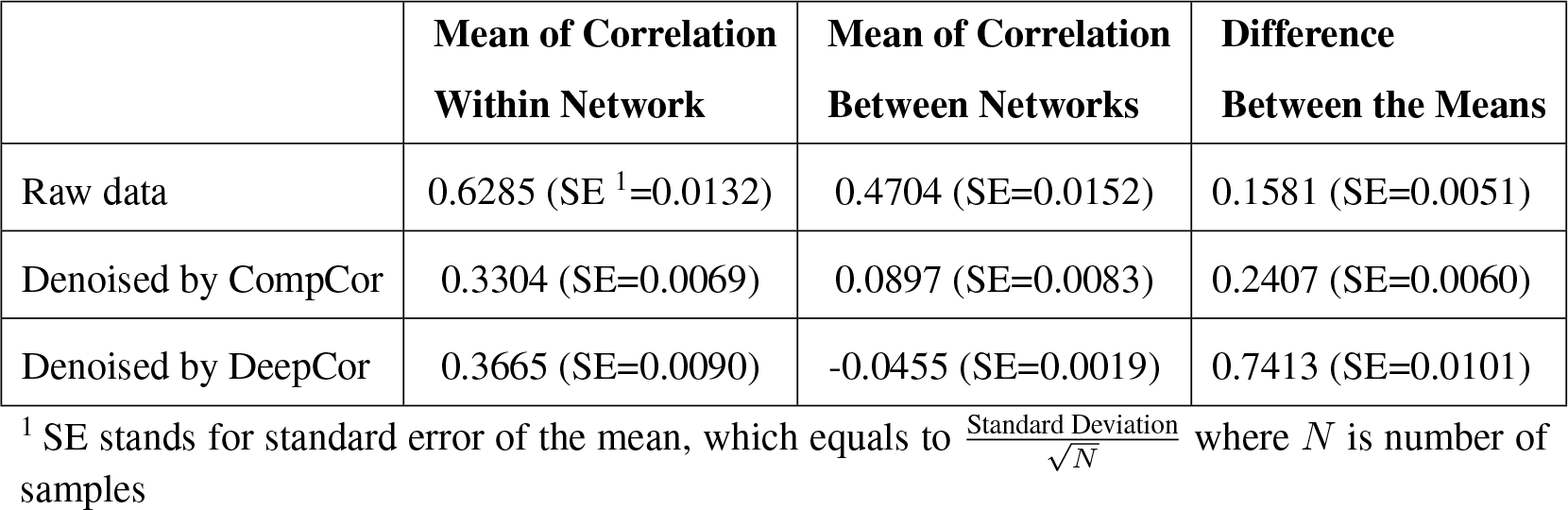
Connectivity Within and Between Networks

Because different fMRI scans can have different numbers of timepoints, the network has to be adapted to match the size of the inputs. To do this, the padding sizes of the encoder and the padding and out-padding sizes of the decoder can be calculated using the algorithm outlined in Figure S1. Due to the number of layers specified (i.e., 4), the number of timepoints needs to be larger than 31 (which would correspond to a 62 seconds long fMRI scan using a 2 seconds TR).

#### Model training

During training, timecourses from the ROI are encoded using both encoders, and reconstructed from their concatenated outputs. Timecourses from the RONI, instead, are only encoded with 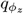, and the features for the signal are set to zero, thus the decoder takes as input the concatenation of the features for the noise and a vector of zeros with the same dimensionality as the features for the signal. As a data augmentation procedure, timecourses in the RONI dataset were multiplied by a scalar *γ* sampled from a Beta distribution (*γ* ∼2∗ *Beta*(4, 4)), with the goal of making the model more robust to variations in the amplitude of the noise.

The loss function is the sum of two terms: a reconstruction loss, and a term comprising the Kullback-Leibler divergences between the encoders’ distributions and the prior, which is set to a standard Gaussian distribution with zero mean and unit standard deviation (as is typical in variational autoencoders, [21]). For the analyses in this article, we used the Adam optimizer with a learning rate of 0.001, *β*_1_ = 0.9, *β*_2_ = 0.999, and *ϵ* = 10^*−*8^.

The model was trained for 50 epochs for the analyses with the simulated dataset using trigonometric functions, 5 epochs for the analyses with the realistic data simulation, and 10 epochs for the application to real data to calculate functional connectivity.

The implementation of the model and the training code is available on Github at this link: https://github.com/sccnlab/DeepCor.git.

#### Data Simulations with Trigonometric Functions

In the first set of simulations, based on trigonometric functions, timecourses in the region of no interest (RONI) were generated with the following Equation 1:

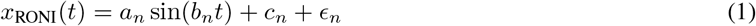

Timecourses in the region of interest (ROI) were generated with the following Equation 2:

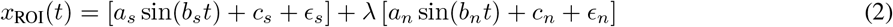

for the case in which signal and noise are combined linearly, and with Equation 3:

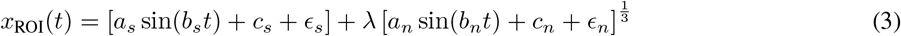

for the case in which signal and noise are combined nonlinearly.

We used the following parameters: *t* = [0, 1, 2, · · ·, 155], *a*_*s*_, *a*_*n*_ ∼*U* (5, 10), *b*_*s*_ ∼ *U* (1, 3), *b*_*n*_ ∼ *U* (0.1, 1), *c*_*s*_, *c*_*s*_ ∼*U* (200, 300), and *ϵ*_*s*_, *ϵ*_*n*_ ∼*U* (−1, 1). All parameters are independently generated for each voxel, except *ϵ*_*s*_ and *ϵ*_*n*_, which are regenerated for every time point. Moreover, *λ* is used to control the standard deviation of the noise proportion, that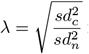for linearly combined cases and 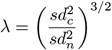 for non-linearly combined cases, where *sd*_*c*_ is the standard deviation we expect the noise proportion to contain and *sd*_*n*_ is the standard deviation the noise proportion contains before transformation. *sd*_*c*_ is constrained to be 0.5, 1.0, and 2.0, respectively for linearly and non-linearly combined conditions. For each trial, a total of 10,000 voxels were simulated for the ROI, and another 10,000 for the RONI.

#### Realistic Data Simulations

To generate more realistic simulations, we employed the fmrisim module of the BrainIAK package [22, 16]. We firstly extract noise parameters from one participant’s fMRI data provided by DataSpace (http://arks.princeton.edu/ark:/88435/dsp01dn39x4181), including the Signal to Noise Ratio (SNR), the Signal to Fluctuation Noise Ratio (SFNR), and the Full Width at Half Maximum (FWHM) [23]. We used a brain mask obtained from the first participant in Study Forrest dataset (https://www.studyforrest.org/) [24] with regions classified by the probability maps with a threshold of 0.5. Utilizing noise parameters derived from empirical fMRI data, we proceeded to generate and combine system and temporal noises, including physiological, drift, autoregressive, and task noise components (Figure 2b), obtaining noise timecourses with a length of 156 timepoints for each voxel in the brain mask. The ground truth of the signal was obtained by simulating responses to two distinct event types, that were observed to occur independently. Each voxel in the ROI was associated with two beta parameters -one for each event type - capturing the amount of response in that voxel to that event type. The ground truth in a voxel in the ROI was then obtained by convolving event timecourses with a hemodynamic response function, and weighting the resulting predictors by the corresponding beta parameters (refer to Figure 2c). Akin to the previously generated simulated datasets, the noise and ground truth signal were summed, yielding the simulated measurements in the ROI. The RONI dataset was obtained with an analogous procedure, but including only the noise (setting the signal to zero). All steps are based on the standard pipeline of the fmrisim module. The resulting data were denoised using both CompCor and DeepCor.

**Fig. 3.**
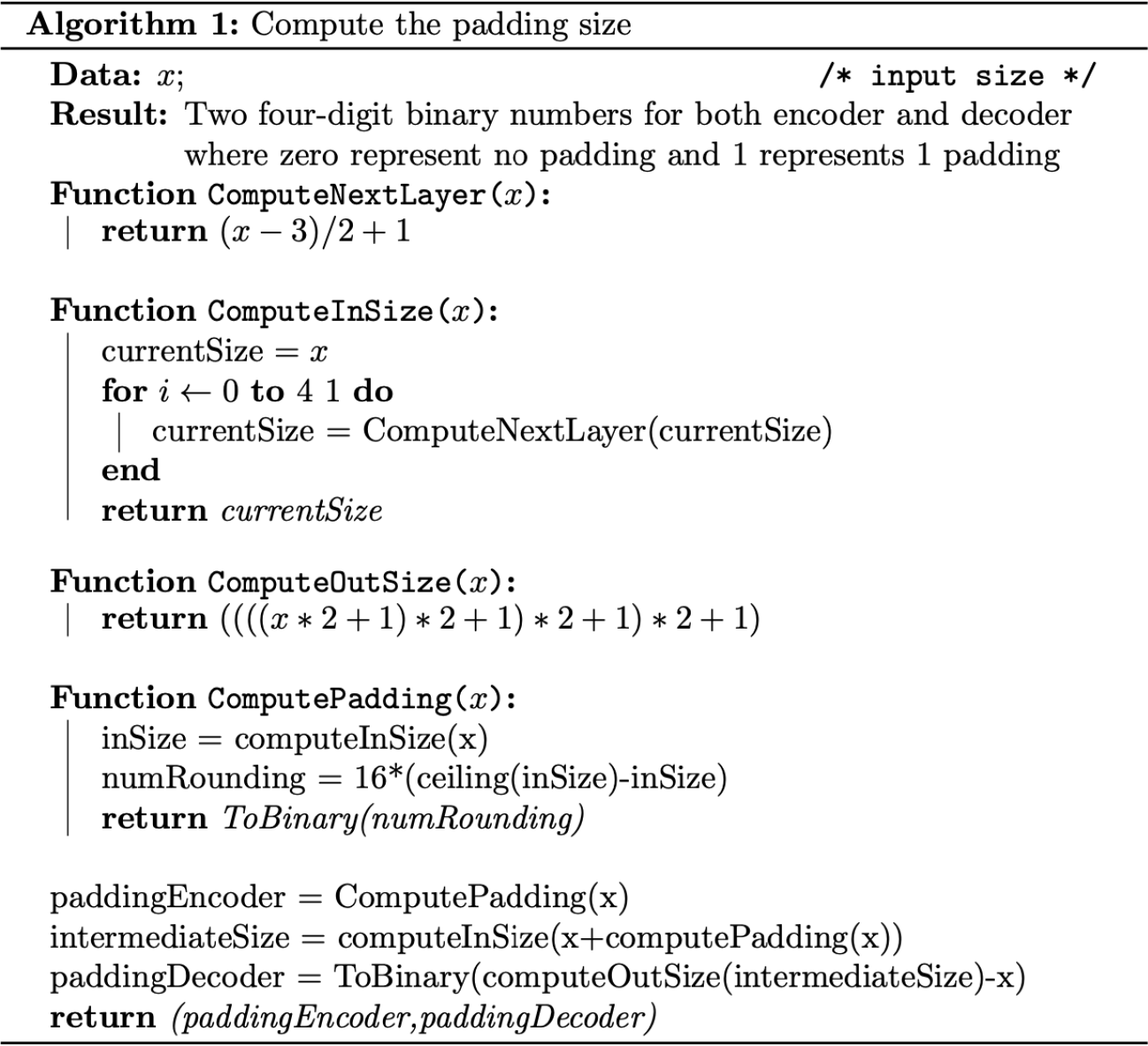

#### Comparison to CompCor

To evaluate the effectiveness of DeepCor, we compared its performance to the performance obtained with a widely used denoising method: CompCor [7]. We denoised the same simulated datasets using both DeepCor and CompCor. For CompCor, we first extracted the top 5 principal components (PCs) from the RONI dataset, and then performed a linear regression using these PCs as predictors, and timecourses in the ROI as target of prediction. Finally, we subtracted the prediction of this model from the original ROI timecourses, obtaining the residuals. The residuals are the part of the ROI timecourses that could not be predicted based on the RONI (noise) timecourses, and thus correspond to the the denoised data.

We assessed the effectiveness of both denoising methods by calculating the R-squared: the square of the correlation between the denoised data and the groudn truth. Since R-squared values violate the normality assumption made by t-tests, we used nonparametric statistical tests. Specifically, Wilcoxon matched pair signed rank test was performed for each simulation on the hypothesis that the mean of R-squared of the sample denoised by DeepCor is greater than the mean of R-squared of the sample denoised by CompCor.

#### fMRI preprocessing

Data were preprocessed with fMRIPrep 22.0.0rc3 (RRID:SCR 016216) [25, 26], which is based on Nipype 1.8.2 (RRID:SCR 002502) [27, 28]. Standard preprocessing steps were included: motion correction, alignment to T1w reference and spatial normalization to MNI152NLin2009cAsym res-2 template at 2mm isotropic resolution. Slice timing correction was not applied. Brain tissue was segmented into tissue classes (gray matter, white matter, cerebro-spinal fluid), nuisance regressors (such as aCompCor components) were calculated as implemented in fMRIprep 22.0.0rc3. The first five time points were discarded for each voxel to remove noises generated before the RF coils warm up and the magnetic field stabilises. No further preprocessing was performed.

#### Application to Functional Connectivity

We randomly selected 200 participants’ fMRI and MRI data from the ABIDE I dataset (http://fcon1000.projects.nitrc.org/) [20], and preprocessed the data following the steps described in the previous paragraph. For each participant, we generated three versions of the data: one without any denoising, one denoised with CompCor, and one denoised with DeepCor (including removal of the global signal). We downloaded masks for the 51 connectivity defined parcels in [19], and their subdivision into 7 networks (https://github.com/ThomasYeoLab/CBIG/blob/master/stable_projects/brain_parcellation/Yeo2011_fcMRI_clustering/1000subjects_reference/Yeo_JNeurophysiol11_SplitLabels/MNI152/Centroid_coordinates/Yeo2011_7Networks_N1000.split_components.FSL_MNI152_2mm.Centroid_RAS.csv. The data was normalized to MNI space, and subsequently denoised. Denoised functional data for different voxels within the same parcel were averaged and processed by a third order Butterworth bandpass filter, removing frequencies lower than 0.01 or higher than 0.1. Finally, we calculated the correlations between the denoised and bandpass-filtered timecourses for each pair of regions, obtaining a connectivity matrix for each version of the data (raw data, CompCor-denoised, and DeepCor-denoised Figure 2g). To assess the statistical significance of differences in functional connectivity between denoising approaches, we first applied the Fisher transformation to each individual connectivity. Next, we averaged the connectivity values for pairs of regions within a same network, and for pairs of regions in different networks. Finally, we computed the difference between the within-network connectivity values and the between-network connectivity values for each participant (see 5). A Wilcoxon matched pair signed rank test was performed over 200 participants, testing whether the differences between the within-network connectivity values and the between-network connectivity values in the sample denoised with DeepCor is significantly larger than in the sample denoised with CompCor.

